# Does host diversity beget microbiome diversity? Effects of zooplankton diversity and composition on environmental and host-associated microbes

**DOI:** 10.64898/2026.09.25.754487

**Authors:** Joshua H. Dominguez, Christopher B. Wall, Margaret Y. Demmel, Jonathan B. Shurin

## Abstract

Microbiomes alter host health and are shaped by transmission within and among host taxa. Microbes can alter host communities, yet whether the diversity of hosts reciprocally shapes microbial communities remains poorly resolved. Diverse host communities may promote microbiome diversity through elevated cross-species microbial spillover or reduce it by diluting conspecific transmission. Here, we manipulated the composition and diversity of pond zooplankton communities to test how host diversity and density alter host-associated and environmental microbiomes. Host diversity had opposing effects on microbiome alpha diversity for different zooplankton host species, increasing microbiome diversity in calanoid copepods while reducing it in *Daphnia*, with no effect on environmental microbes. Host diversity amplified cross-species microbial spillover for subordinate competitors (calanoid copepods) and diluted conspecific transmission for the dominant competitor (*Daphnia*). These contrasting responses reflected shifts in indicator microbes associated with each host taxon. Increasing host turnover generated higher microbiome turnover, demonstrating that microbiome assembly depends on host community composition. Our results indicate that contact with both conspecifics and heterospecifics alters animal microbiomes, and that host diversity can either promote the spillover or dilute the transmission of microbes in different zooplankton with potential effects on host health and ecosystems.

## 1. INTRODUCTION

Microbes (bacteria and archaea) shape the ecology and evolution of eukaryotic hosts, and microbial diversity can alter host health and ecosystem functioning (Laforest-Lapointe et al. 2017; Dickey et al. 2025; Corral López et al. 2026). The structure of host-associated microbiomes varies within and among host taxa (Brooks et al. 2016), across space (Suzzi et al. 2023; Härer & Rennison 2023; Wall & Perreault et al. 2025), and over time (McCauley et al. 2020; Marsh et al. 2024). Consequently, unraveling the factors that shape microbiomes is essential to understanding host health, ecological interactions, and ecosystem function. Horizontal transmission of microbes among co-occurring hosts is one process that structures microbiomes (Burns et al. 2017; Moeller et al. 2017; Sarkar et al. 2020). Hosts and their associated microbiomes are linked through networks of interactions that can enable the coexistence of hosts and influence population dynamics (Abbott et al. 2021; Meyer et al. 2022), though these effects are often cryptic and underappreciated.

Dispersal among habitat patches can alter local diversity and community turnover by introducing new species or reshuffling the relative abundance of residents (Mouquet & Loreau 2003; Leibold et al. 2004). In microbiomes, hosts act as habitat patches connected by microbial dispersal through direct interactions or via a shared environmental pool of microbes (Kembel et al. 2012; Mihaljevic 2012; Miller et al. 2018). Therefore, interactions among hosts can facilitate microbial transmission that reshapes microbiome composition and diversity. For example, microbiomes are more similar within animal social groups (Sarkar et al. 2020; Rose et al. 2023; Valles-Colomer et al. 2023; Sarkar et al. 2024), indicating that microbiomes are shaped by contact among hosts. Fish have more diverse microbiomes when reared in groups than in solitary conditions, consistent with enhanced microbial transmission among individual hosts (Burns et al. 2017). These findings suggest that increasing host density promotes contact and microbial transmission among individuals, potentially increasing microbiome diversity in hosts and the environment through repeated colonization by host-specialized microbial taxa.

In natural communities, hosts interact with multiple species that serve as reservoirs of microbial diversity (Brooks et al. 2016; Mallott & Amato 2021). Indeed, the diversity of plant neighborhoods can alter the composition (Meyer et al. 2022; Meyer et al. 2026) and amplify the diversity (Lu et al. 2025) of plant phyllosphere microbiomes, suggesting that interactions with other species reshape microbiomes. However, cross-species exchange of microbes may be prevented by host selection or competition with resident taxa. Manipulations of microbial transmission among conspecific and heterospecific hosts produce divergent microbial communities, with conspecific transmission reinforcing host-specialized microbes and heterospecific transmission favoring generalist microbiomes (Meyer et al. 2023). In addition, environmental microbial communities can be influenced by the composition of the host community (Miller et al. 2018; Macke et al. 2020). However, few studies test how the diversity of hosts influences either their associated microbiomes or free-living microbial assemblages.

Host diversity may affect microbial diversity in the environment and in hosts through several potential mechanisms. If host taxa share microbial symbionts, then hosts in diverse communities may experience greater horizontal transmission from heterospecifics and therefore have more diverse microbiomes. This mechanism depends on the ability of microbes to successfully colonize multiple host taxa. Spillover of microbes from different host taxa may also enrich environmental microbial assemblages. Alternatively, if different macroscopic taxa cannot host the same symbionts, then there may be little potential for cross-species sharing of microbes. In this case, competition among diverse hosts may reduce the density of each host species and therefore reduce the potential for horizontal transfer among conspecifics. By this mechanism, analogous to the “dilution effect” in disease ecology (Schmidt & Ostfeld 2001; Keesing & Ostfeld 2021), increasing host diversity may reduce microbiome diversity within hosts by decreasing opportunities for conspecific microbial transmission. The diversity of plant and animal assemblages may therefore either promote or suppress microbial diversity depending on its effect on host density and the potential for cross-species sharing of microbes.

Here, we experimentally manipulated the diversity and composition of pond crustacean zooplankton communities to test how host community structure alters microbiomes in hosts and the environment. Freshwater zooplankton, particularly *Daphnia*, are an emerging model system for studying host-microbe interactions given their critical roles in food webs and sensitivity to microbiome composition (Sison-Mangus et al. 2015; Akbar et al. 2022). Using replicated aquatic mesocosms, we assembled zooplankton communities seeded with one, two, or three taxa (*Daphnia*, *Calanoida*, and *Cyclopoida;* Fig 1C) to generate a gradient of host Shannon diversity. We tracked the diversity and composition of microbes in zooplankton hosts and water at three time points over a six-week span. We tested two hypotheses of how host diversity reshapes host-associated microbiome diversity. Increasing host diversity may enhance cross-species exchange of microbes, amplifying microbiome diversity (Fig 1B). Alternatively, increasing host diversity may dilute transmission among conspecific hosts with shared symbionts, lowering microbiome diversity (Fig 1B). The direction and magnitude of these effects may depend on the density of hosts – whether conspecific or heterospecific – in the community. We use indicator species analysis to identify bacterial taxa associated with each of the three hosts and ask how their prevalence is shaped by the composition of the zooplankton community. Our study tests how variation in host communities reshapes microbiomes in hosts and ecosystems.

**Figure 1.**
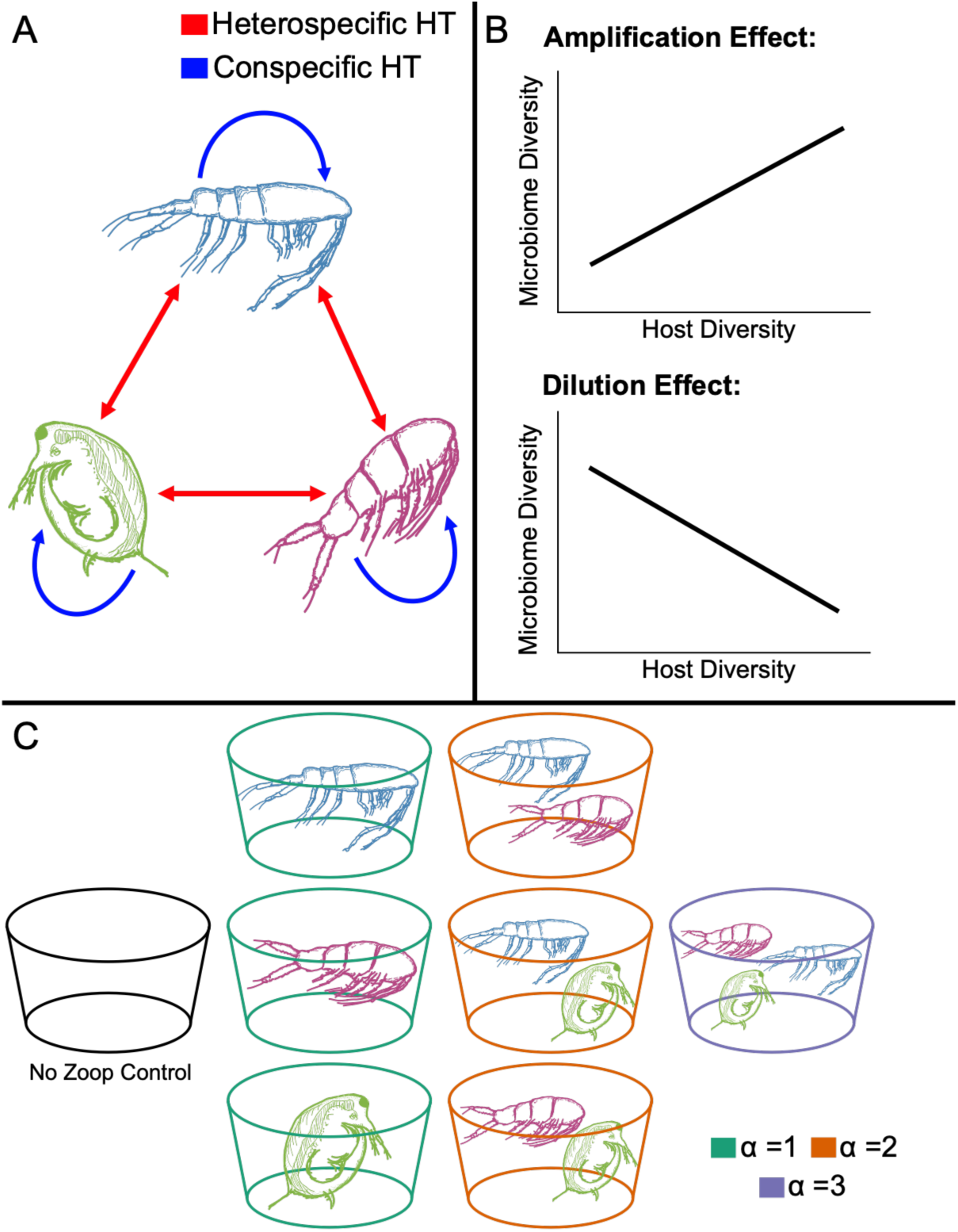
**(A)** Conceptual diagram of conspecific (blue) and heterospecific (red) microbial horizontal transmission among hosts in a community. Variation in host community structure (diversity and density) can shape the strength of conspecific and heterospecific microbial transmission, with downstream effects on host-associated and environmental microbial diversity. **(B)** Hypotheses for how host diversity alters microbiome diversity. If increasing host diversity enhances opportunities for microbial transmission among hosts, microbiome diversity may increase (amplification). Alternatively, if increasing host diversity reduces conspecific transmission or if heterospecific microbes fail to establish, microbiome diversity may decline (dilution effect). **(C)** Schematic of the host diversity experiment. Mesocosms were stocked with zooplankton communities dominated by one (α = 1), two (α = 2), or three (α = 3) focal taxa (*Daphnia* – green, *Calanoida* – blue, *Cyclopoida* – magenta).

## 2. METHODS

### 2.1 Experimental Design

In the summer of 2022 (July – August), we established a gradient of host diversity and density that varied through time in 1000 L aquatic mesocosms at the Sierra Nevada Aquatic Research Laboratory (Fig 1C). We used three focal crustacean zooplankton taxa: *Daphnia* (*Daphnia* spp.), cyclopoid copepods (*Cyclopoida* spp.), and calanoid copepods (*Calanoida* spp.). Zooplankton were collected from Eastern Brook Lake (37.43153°N, 118.74262°W) using a 64-μm mesh zooplankton net and transported live to the laboratory in thermoses. Zooplankton taxa were isolated under a dissecting scope (Leica Biosystems) and transferred to 1-L Nalgene bottles. We seeded mesocosms to create three discrete richness treatments consisting of control tanks with no zooplankton, single taxon tanks, tanks with two taxa in pairwise, and tanks with three taxa (n = 8 community treatments, in three replicate blocks, totaling 24 tanks; Fig 1B). Low-level cross-colonization among tanks and natural community dynamics drove assemblages to diverge from initial treatment conditions over time, generating a continuous gradient of host diversity and density. Zooplankton diversity was related to the initial experimental treatment (Fig 2). Consequently, we evaluated host community structure as continuous variables (i.e. Shannon diversity and density) rather than categorical treatments. We stocked mesocosms daily for two weeks to ensure sufficient zooplankton densities and then left tanks unmanipulated (see Supporting Methods for details).

**Figure 2.**
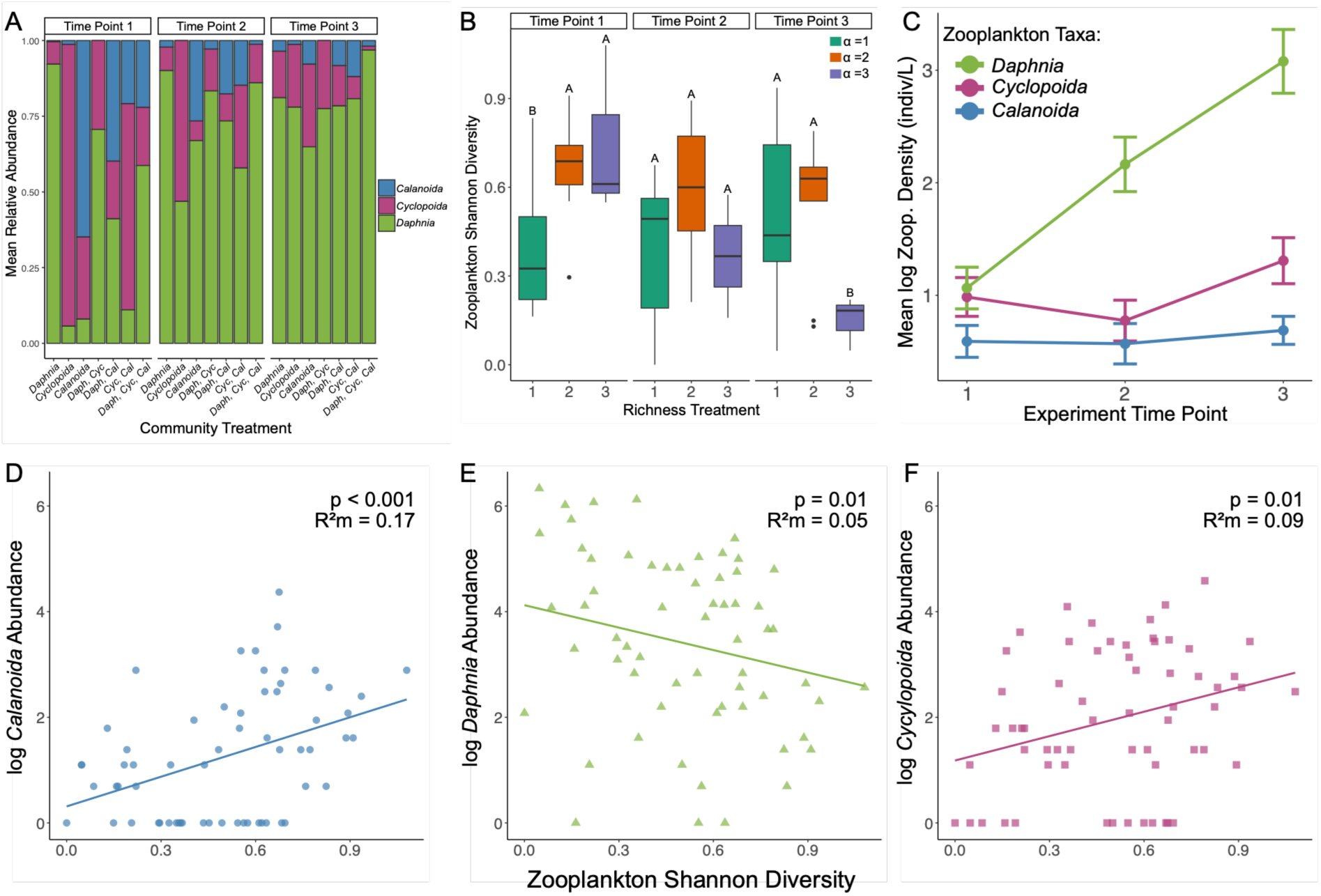
Zooplankton community structure. **(A)** Barplot of focal taxa mean relative abundances in community treatments across time points. **(B)** Boxplots of zooplankton Shannon diversity in different richness treatments (1-3 taxa) over time. Richness treatments with significantly different Shannon diversity within time points based on post-hoc tests (p < 0.05) are marked by letters. **(C)** Mean log transformed densities (indiv/L) of zooplankton taxa across tanks over time. Points are colored by focal taxa (*Calanoida* – blue*, Daphnia* – green, *Cyclopoida* – magenta). **(D-F)** Zooplankton Shannon diversity as a function of focal taxa abundances.

### 2.2 Zooplankton and Microbiome Sampling

We sampled zooplankton and tank water for host community and microbiome analyses at three time points 14, 35, and 44 days after zooplankton stocking ceased. For bacterioplankton, surface water was pre-filtered through a 64-μm mesh and collected in sterile, acid washed (10% HCl) 300 mL Nalgene bottles. We immediately filtered water samples through 0.2-μm filters (Supor membrane, Pall Corporation, New York, NY, USA) and stored them at −80°C.

Zooplankton were sampled from each mesocosm using 1-L vertical water samplers specific to each community treatment (4-L sampled per tank). For community analysis, zooplankton were filtered (>64-μm) and preserved in 70% ethanol. Zooplankton samples were counted under a dissecting scope (Leica Biosystems) and identified to focal taxa levels (*Daphnia, Calanoida, Cyclopoida*). Taxa and community-level densities were calculated as the number of individuals per liter of tank water sampled.

To examine host microbiomes, living zooplankton from tanks at each time point were transferred to sterile microcentrifuge tubes filled with 100% molecular grade ethanol and stored at −80°C for downstream DNA extraction following Wall & Perreault et al. 2025.

### 2.3 DNA Extraction, Sequencing, and Bioinformatics

Zooplankton preserved at −80°C for microbiome analysis were separated by taxa into DNA extraction tubes using a dissecting scope under sterile conditions (flamed dental tools and sterile petri dishes). Zooplankton microbiomes included the whole body, representing internal and external associated bacteria and archaea. We pooled between 1 and 32 individuals per extraction tube based on previous analysis (Demmel & Wall et al. 2025; Wall & Perreault et al. 2025). Up to four replicates per focal taxon were sampled within tanks at each time point. A total of 197 zooplankton samples were included in downstream processing. We extracted genomic DNA from all samples and sent aliquots to the Argonne National Laboratory (ANL) for library preparation and 250 × 250 bp paired-end Illumina MiSeq sequencing on the V4 region of the 16S-rRNA gene (Walters et al. 2016) (Supporting Methods for details).

Illumina MiSeq reads were demultiplexed and fastq files were run through the DADA2 pipeline to identify ASVs (Callahan et al. 2016) (Supporting Methods for details). Our final dataset consisted of 243 samples (92 *Daphnia*, 56 *Calanoida,* 41 *Cyclopoida,* and 54 bacterioplankton) with 5,497 unique ASVs. Down-stream analyses were performed using R Version 4.4.2 (R Core Team, 2025).

### 2.4 Zooplankton Community Analysis

We analyzed zooplankton Shannon diversity, density, and community composition over time. We tested the interactive effects of initial richness treatment (1-3 taxa) and time point on Shannon diversity using a Gamma GLMM with tank as a random effect using ‘glmmTMB’ (Brooks et al. 2017). Zooplankton density was modeled using a negative binomial GLMM with host taxa and time as fixed effects. Pairwise differences among richness treatments and zooplankton taxa within each time point were assessed using Tukey-adjusted estimated marginal means (*emmeans*; Lenth & Piaskowski 2025). Temporal shifts in community composition were evaluated using PERMANOVA (Bray-Curtis distances; *adonis2* in ‘vegan’; Oksanen et al. 2007) with permutations constrained within tanks to account for repeated measures. We fit two separate models evaluating richness and community treatments because richness was nested within community treatment and fit additional PERMANOVA models within each time point.

### 2.5 Microbiome Composition

We analyzed the composition of zooplankton and water microbiomes using Bray-Curtis (Hellinger-transformed ASV count matrices) and weighted UniFrac distances (*distance()* function in ‘phyloseq’; McMurdie & Holmes 2013). We used PERMANOVA to test for effects of sample type, time, and host community structure on microbiome composition. Two complementary models were fit: the first testing the effects of sample type and time, and the second testing the effects of realized host Shannon diversity and density, including their interactions with sample type. Permutations were constrained within tanks to account for repeated measures. Differentially abundant taxa among sample types were identified using ANCOM-BC at the Class level (Lin & Peddada 2020). Differences in microbiome alpha diversity among sample types were evaluated using Gamma GLMMs with tank as a random effect.

### 2.6 Effects of Host Density and Diversity on Microbiome Alpha Diversity

We tested the effects of zooplankton Shannon diversity and host density on three indices of microbiome alpha-diversity (Shannon Diversity, ASV Richness, and Pielou’s Evenness Index) using GLMMs. Because host Shannon diversity covaried with the densities of individual host taxa (Fig 2), we evaluated the effects of host diversity and density in separate models. For each host, we fit a diversity model including host Shannon diversity and a density model including conspecific and heterospecific host densities. For environmental microbiomes, the density model included total host density of all zooplankton species. We compared models using AICc to assess whether diversity or the density better explained variation in microbiome diversity. Time point and tank were included as random intercepts in all models. Shannon diversity was modeled with a Gamma distribution, ASV richness with a negative binomial distribution, and Pielou’s evenness with a beta distribution. We report marginal (R^2^m; variance explained by fixed effects) and conditional (R^2^c; variance explained by fixed and random effects) R^2^ values for mixed effects models.

### 2.7 Microbial Indicator Species Analysis

We tested how host community structure influences host-specialized microbes by modeling the relative abundance of microbial indicators as a function of the density of the host they are indicators of (conspecifics) and the combined density of other hosts (heterospecifics). We aggregated ASVs to the Family level and performed indicator analysis (*multipatt()* in ‘indicspecies’; De Cáceres 2013) to identify taxa associated with individual hosts (host indicators), multiple hosts (zooplankton generalists), or combinations that included environmental samples (water-associated taxa). We summed the relative abundances of indicators for each corresponding host. Next, we fit generalized linear mixed models predicting the summed relative abundance of host-indicator microbes as a function of conspecific and heterospecific host densities for each host taxa. Models were fit using a beta distribution with a logit-link and indicator taxa relative abundances were rescaled to the open unit interval to account for exact 0 and 1 values in our data (Smithson & Verkuilen 2006). Time point and tank were treated as random effects in all models.

### 2.8 Effects of Zooplankton Composition on Microbial Turnover

To test whether variation in host community turnover shapes microbiome turnover through time, we modeled microbiome β-diversity as a function of host β-diversity (Bray-Curtis dissimilarities). Microbiome dissimilarities were calculated within sample types, and comparisons from the same tank and time point were excluded. We fit Bayesian GLMMs with a Beta distribution and logit link (‘brms’ R package; Bürkner 2017). Fixed effects included time between samples (days, z-standardized) and host β-diversity (z-standardized) with interactions by sample type. Non-independence among comparisons was modeled using multi-membership (mm) random effects for sample identity and tank: mm(sample *i*, sample *j*) and mm(tank *i*, tank *j*) with equal weights. Variance inflation factors were low (VIF ≤ 2.54), indicating weak collinearity between time and host beta-diversity. Effects were considered supported when 95% credible intervals excluded zero.

## 3. RESULTS

### 3.1 Zooplankton Community Structure

Zooplankton Shannon diversity differed among richness treatments early in the experiment but converged over time (richness × time point interaction; Fig 2B; Table S1; Table S2), indicating a breakdown of the initial categorical design into continuous variation of host diversity. Host densities (i.e. abundances) did not differ among taxa at the initial time point (Tukey-adjusted P > 0.05; Table S4) but varied over time (host × time interaction; Table S3; Fig 2C). By the second time point, *Daphnia* exhibited significantly higher densities than both copepods (Tukey-adjusted P ≤ 0.001). In the final time point, all zooplankton taxa differed in densities, with *Calanoida* being the least abundant (Tukey-adjusted P < 0.01; Fig 2C). Overall, *Daphnia* increased over time, whereas copepods remained relatively stable. As a result, host Shannon diversity covaried with host densities, with *Daphnia* dominating low-diversity communities and copepods dominating high-diversity communities (Figs 2D-F).

Zooplankton community composition was shaped by both community treatment (PERMANOVA; R^2^ = 0.26; P = 0.001; Table S5) and time (R^2^ = 0.16; P = 0.001; Table S5). However, community treatment effects weakened over time (Fig S1), indicating divergence from discrete community treatments. In contrast, richness treatment explained comparatively little variation in composition (R^2^ = 0.04; P = 0.001; Table S5). Together, these results show that the experiment generated a continuous gradient of host diversity, density, and community composition over time due to invasion of species in treatments outside where they were initially introduced. However, the tanks varied in zooplankton density, composition, and diversity due to the initial treatments. Consequently, we analyzed the effects of host community structure as continuous predictors over time.

### 3.2 Microbiome Composition

Microbiome composition differed among sample types and time points (PERMANOVA; Table S6; Figs S2–S3). Differences were greatest between hosts and the environment, and *Daphnia* microbiomes were distinct from copepods (Fig S2). Differential abundance analyses revealed broad taxonomic partitioning among sample types at the Class level (Fig S2C).

Host community structure had comparatively weak effects on microbiome composition. Host Shannon diversity explained little variation (sample type x host Shannon; R^2^ = 0.01, P = 0.044; Table S7), whereas total host density had modest but significant effects (sample type x total host density; Bray-Curtis: R^2^ = 0.02, P = 0.001; Table S7). Overall, microbiome composition was primarily shaped by host identity, with host diversity and density playing secondary roles.

### 3.3 Effects of zooplankton community structure on microbiome alpha-diversity

Zooplankton Shannon diversity had taxa-specific effects on host microbiome alpha diversity but did not influence the diversity of environmental communities (Fig 3; Table 1). Host Shannon diversity increased microbial Shannon diversity in calanoids (R^2^m = 0.17; R^2^c = 0.17; P < 0.001; Fig 3A) but decreased Shannon diversity in *Daphnia* (R^2^m = 0.08; R^2^c = 0.54; P = 0.003; Fig 3A). In contrast, host diversity had no detectable effects on the Shannon diversity of cyclopoid copepod (P > 0.05; Fig 3A) or water microbiomes (P > 0.05; Fig 3D). Patterns for microbial richness and evenness (Pielou) were broadly consistent with Shannon diversity but varied in strength among taxa (Table S8). In calanoids, host diversity drove positive effects on evenness (P = 0.002) and marginal positive effects on richness (P = 0.073). In Daphnia, host diversity consistently reduced richness and evenness (P < 0.05; Table S8). In cyclopoids and water microbiomes, host diversity had no effects across all metrics of microbial alpha-diversity (Fig 3; Table S8).

**Figure 3.**
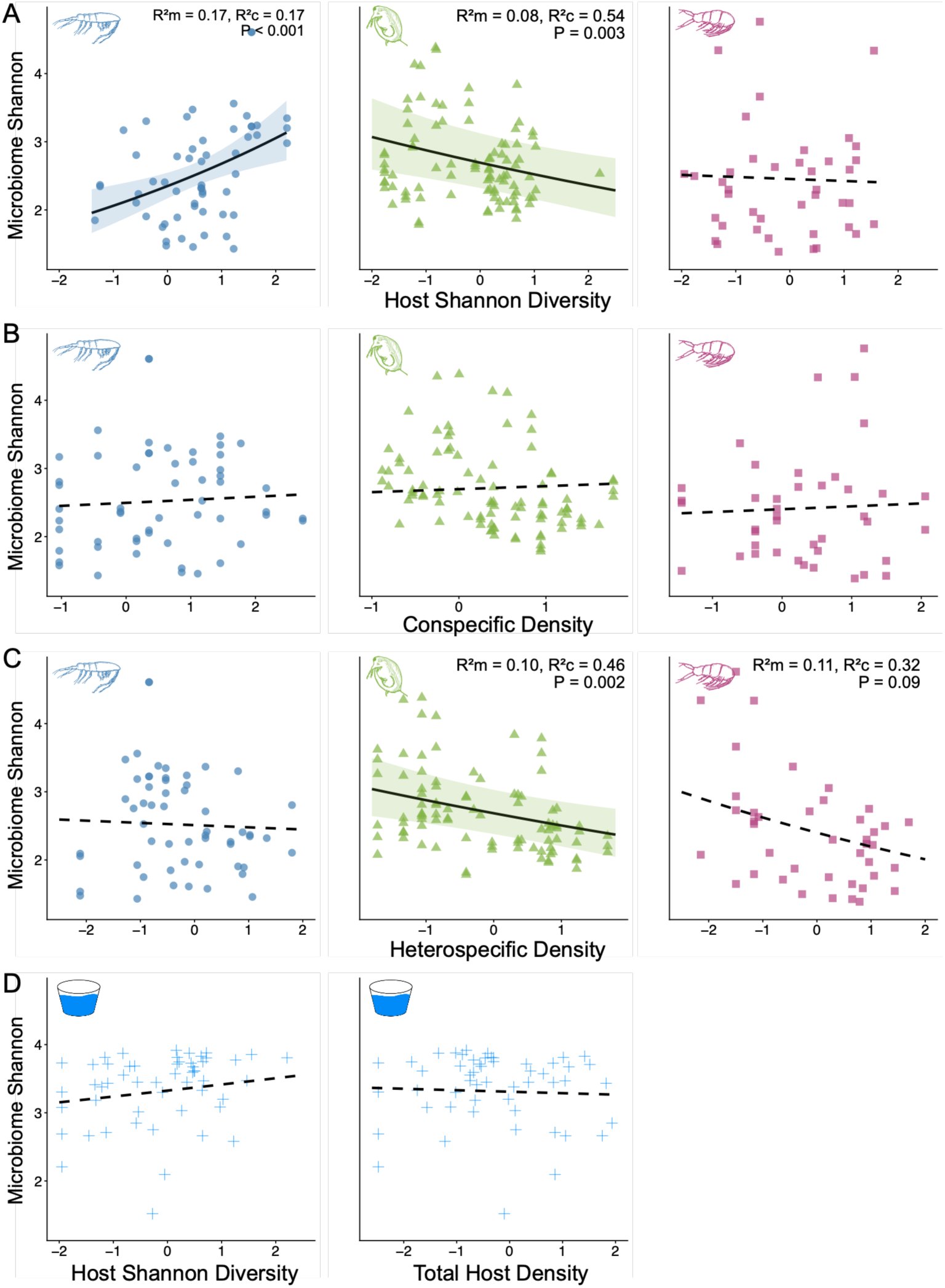
Effects of host community structure on microbiome Shannon diversity. Panels show effects of **(A)** host Shannon diversity, **(B)** conspecific density, and **(C)** heterospecific density on host microbiomes. Panel **(D)** depicts the effects of host Shannon diversity and total host density on water microbiomes. Shape and color denote sample type (*Daphnia* – green triangles; *Calanoida* – blue circles; *Cyclopoida* – magenta squares; Water – blue crosses). Marginal and conditional R^2^ and corresponding P-values are reported for significant predictors (solid lines).

**Table 1.**
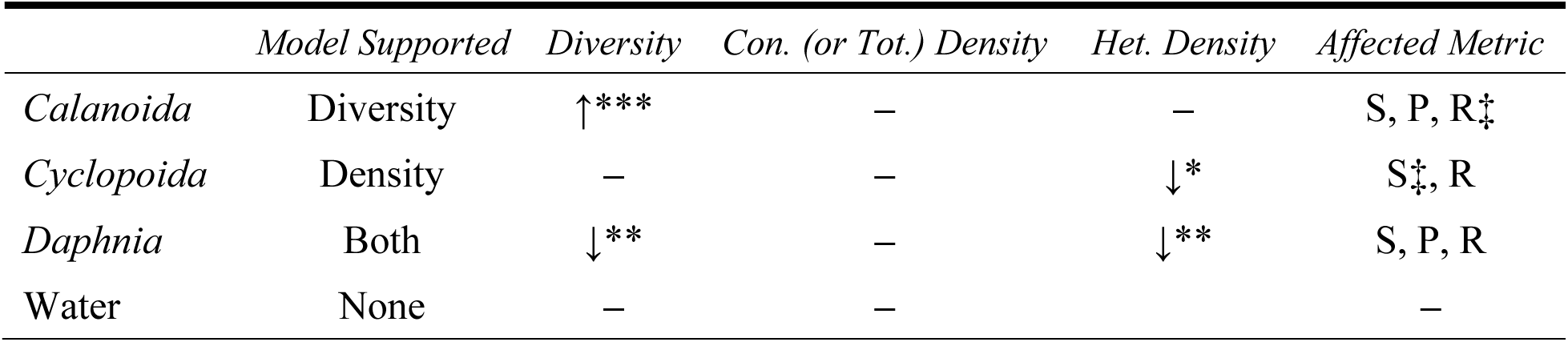
Direction and relative support of host diversity and density effects on microbiome alpha diversity. Arrows indicate the direction of significant effects (↑ positive, ↓ negative), and dashes indicate no effect. Asterisks denote significance levels (* P < 0.05, ** P < 0.01, *** P < 0.001). Affected metrics are abbreviated as S (Shannon diversity), P (Pielou’s evenness), and R (richness), with ‡ indicating marginal support (P < 0.1) for a given metric. Model support is based on ΔAICc, where values < 2 indicate similar support among models. Conspecific density refers to conspecific host density for zooplankton and total host density for water samples. Full model results are provided in Table S8.

Conspecific density did not significantly influence microbiome alpha diversity across taxa (Fig 3B). However, heterospecific density significantly reduced cyclopoid microbiome richness (P = 0.001; Table S8) and had marginal negative effects on Shannon diversity (Fig 3C, p = 0.09). Increasing heterospecific density consistently decreased Shannon diversity, richness, and evenness in *Daphnia* microbiomes (P < 0.05). No density effects were detected for calanoids or bacterioplankton.

Model comparisons (AICc) supported these patterns. Host diversity was a consistent predictor of calanoid microbiome diversity, while host models with host density performed similarly or better for *Cyclopoida* and *Daphnia* (Table 1; Table S8). Neither zooplankton diversity nor density explained variation in bacterioplankton alpha diversity.

### 3.4 Effects of Host Density on Indicator Species Abundance

Host-specialized microbial indicators comprised a small subset of microbial families identified (Fig 5A) yet accounted for a large proportion of host microbiomes, while water-associated taxa dominated bacterioplankton communities (Fig 5B). Host-associated indicator taxa reached their highest relative abundances within their focal hosts, particularly in *Daphnia* and *Calanoida*, while also occurring but less abundant in other hosts (Table S9, Fig 5B). Several taxa showed strong host affinity, including *Comamonadaceae* in calanoids (mean relative abundance 57%) and *Aeromonadaceae* in cyclopoids (41%). *Daphnia* indicators were more evenly distributed (e.g., Candidatus Hepatincola 25%, Flavobacteriaceae 14%; Table S9). In contrast, water-associated taxa were more numerous but rare (Table S9), and generalist microbes comprised small proportions across host microbiomes (Fig 5B).

Host density shaped the relative abundance of microbial indicators in *Calanoida* and *Daphnia* (Fig 4). In *Calanoida*, the relative abundance of indicator microbes exhibited marginal increases with conspecific density (P = 0.08; Table S10). However, *Calanoida* indicator abundance significantly declined as heterospecifics became denser (P < 0.001; Fig 4C; Table S10). Similarly, *Daphnia* indicator abundance marginally increased with conspecific density (P = 0.06; Fig 4C), however, heterospecific density had no effect (Table S10). *Cyclopoida* indicator abundance was not sensitive to changes in conspecific or heterospecific density (Table S10). In summary, the abundances of microbial taxa closely associated with specific hosts are sensitive to the composition of the host community.

**Figure 4.**
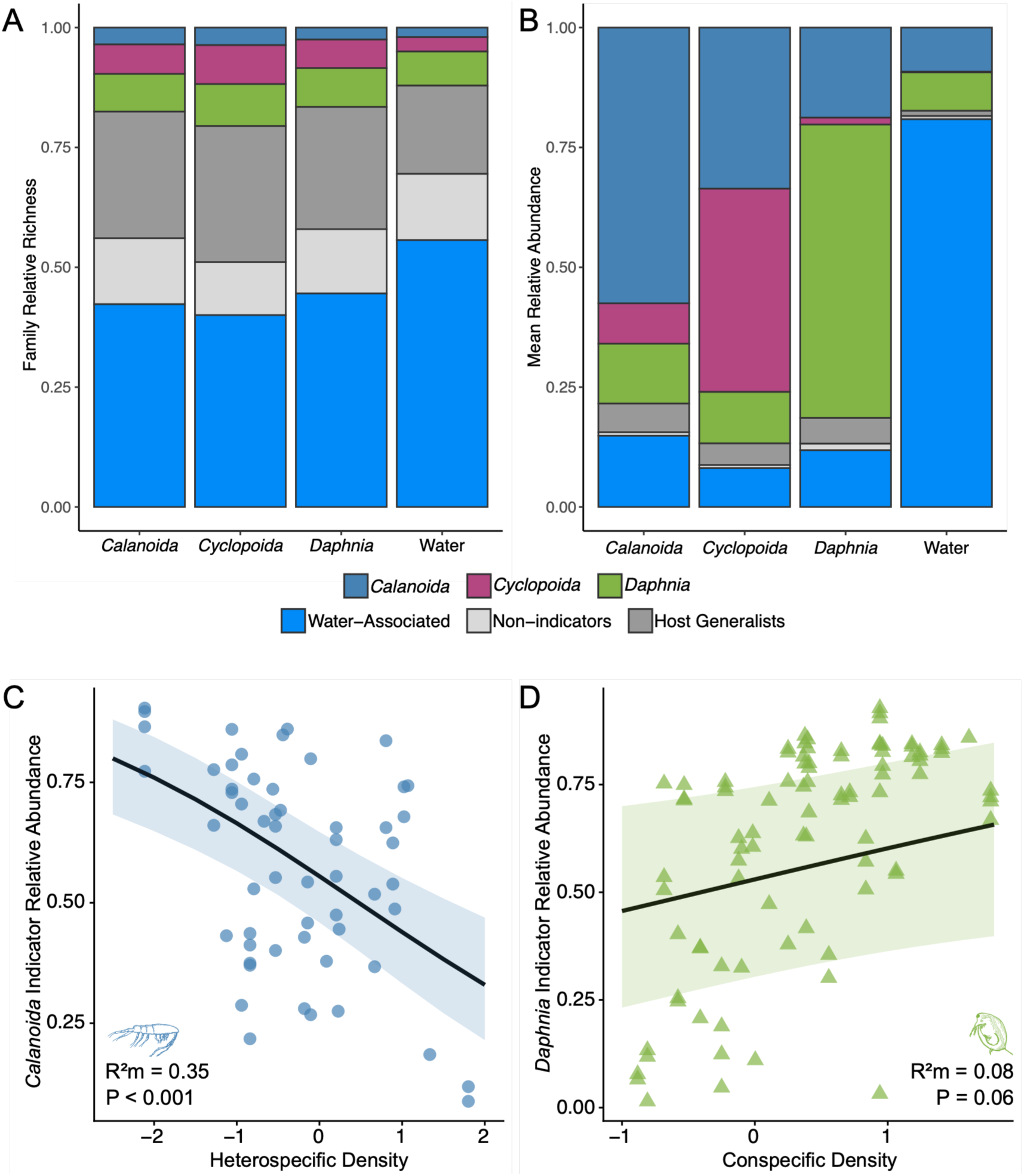
Indicator species analysis. **(A)** Mean proportion of microbial families belonging to each indicator category across host and environmental microbiomes. **(B)** Mean relative abundances of focal host-associated indicators, host generalists, non-indicators, and water-associated microbial families across sample types. **(B)** Marginal effects plot showing the negative relationship between heterospecific host abundance (*Daphnia* + *Cyclopoida*) and the relative abundance of *Calanoida* indicator families within calanoid microbiomes. **(C)** Marginal effects plot showing the positive relationship between conspecific host abundance (*Daphnia*) and the relative abundance of *Daphnia* indicator families within *Daphnia* microbiomes. Lines represent predicted values from regression models, and shaded ribbons represent 95% confidence intervals.

### 3.5 Effects of Zooplankton Composition on Microbial Turnover

Zooplankton turnover predicted microbial turnover in two of the three host taxa but did not influence environmental microbial communities (Fig 5A, Table S11). Host turnover had positive effects on microbial turnover in *Calanoida (*β = 0.09, 95% CrI [0.07, 0.12]) and *Daphnia* (β = 0.08, 95% CrI [0.06, 0.09]), whereas effects in *Cyclopoida* were weak (β = 0.03, 95% CrI [0.00, 0.06]). Turnover in water microbiomes was not sensitive to zooplankton turnover (β = −0.01, 95% CrI [-0.04, 0.02]). Time strongly predicted microbial turnover across all sample types (Fig 5B; Table S11), with the largest effect in water microbes (β = 0.30, 95% CrI [0.28, 0.33]) compared to zooplankton microbiomes (*Calanoida*: β = 0.13, 95% CrI [0.11, 0.15]; *Cyclopoida*: β = 0.14, 95% CrI [0.11, 0.18]; *Daphnia*: β = 0.24, 95% CrI [0.23, 0.26]).

**Figure 5.**
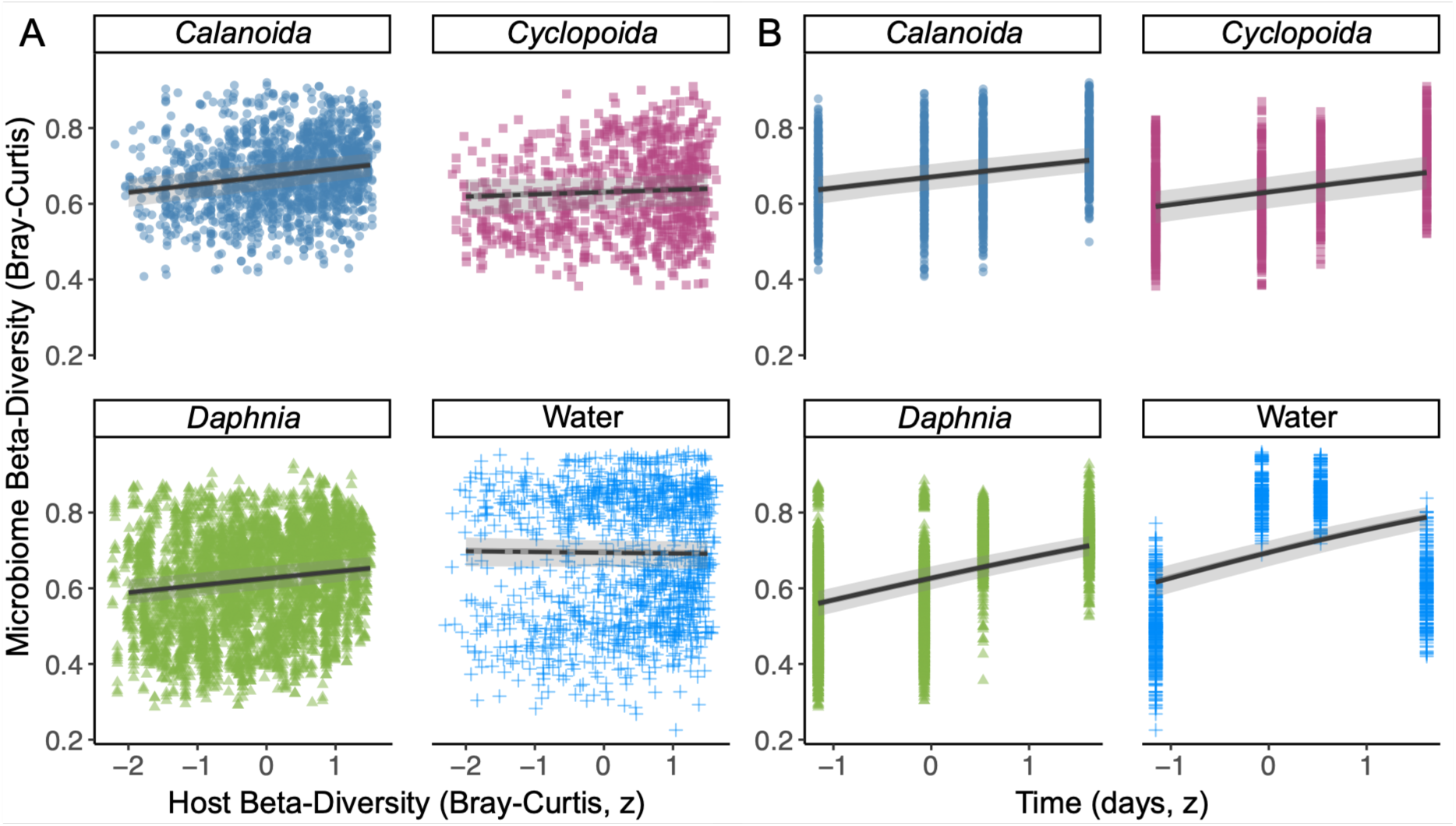
Effects of host β-diversity (Bray-Curtis) and time (days between sampling) on microbiome β-diversity. **(A)** Marginal effects plot showing the relationship between host β-diversity and microbiome β-diversity across sample types. **(B)** Marginal effects plot showing the relationship time difference in days and microbiome β-diversity across sample types. Credible relationships (95% CrI excludes zero) are solid lines, dashed lines indicate effects whose intervals overlap zero.

## 4. DISCUSSION

Theoretical and empirical studies show that horizontal transmission of microbes among hosts can alter microbial communities, yet empirical evidence for feedbacks between host and microbial diversity is rare. Our field experiment demonstrates that changes in host community structure alter the diversity, composition, and turnover of host-associated microbiomes but have little effect on environmental microbes. Zooplankton diversity had host-specific effects on microbiome diversity. Of the three zooplankton in our experiment, *Daphnia’s* microbiome diversity increased with host diversity, while that of *Calanoida* declined and *Cyclopoida* showed no change. These divergent outcomes were related to the association between host density and diversity. *Daphnia,* the dominant host, was most abundant in low diversity communities, while the two copepods were most abundant when zooplankton diversity was highest and *Daphnia* density was lowest. The effect of host diversity was therefore mediated by its effects on host density. These results suggest that transmission within host species is more important than transmission between different host species to microbiome diversity.

### 4.1 Host community structure shapes host, not environmental, microbiomes

Our experiment found that host community structure (the abundance, composition, and diversity of hosts) influenced the diversity and composition of host-associated microbiomes but not environmental microbes. Changes in host communities are expected to alter environmental microbial communities by modifying the transmission of microbes among hosts via the external environment (Miller et al. 2018). For example, manipulative plant experiments show that plant species richness and plant identity can alter soil microbial community composition and function (Zak et al. 2003; Lange et al. 2015; Burns et al. 2015; Chen et al. 2019). In contrast, environmental microbiomes in our aquatic system were insensitive to zooplankton host Shannon diversity, beta-diversity, and density across all tested metrics of microbial diversity and composition (Fig 3, Fig 5A). Microbes in soils may be more locally constrained to the neighborhood of host plants than those in pelagic systems where flow and turbulence mix plankton. Instead, microbiome responses to host community structure were confined to hosts, with *Calanoida* and *Daphnia* microbiomes consistently tracking variation in host communities (Figs 3–5). This result indicates that microbes associated with zooplankton make up a small part of the free-living community, and microbes are transmitted across species even though these taxa are rarely found in the environment.

A strong affinity of microbes for specific animal hosts likely limits their persistence in external environments. Horizontal transmission requires host-associated microbes to persist in the environment long enough to colonize new hosts, yet microbial symbionts may experience rapid mortality or competitive exclusion once released from host reservoirs. Our results suggest that transmission among hosts may occur through undetectable, transient environmental exposure rather than through sustained restructuring of environmental microbial communities. Many microbes associated with animals are highly adapted to their hosts and are rare in environmental communities across a range of systems (Thompson et al. 2017; Suzzi et al. 2023; Dominguez et al. 2026). Our data reflect this dynamic as host and environmental microbes were consistently differentiated (Fig S2A; Fig S2B; Table S2) and shared few ASVs (Fig S3A). Zooplankton-associated microbes are consistently distinct from water samples (Wall & Perreault et al. 2025; Demmel & Wall et al. 2025; & Li et al. 2026), suggesting that a portion of zooplankton-associated microbes are either rare in the environment or are poorly adapted to persist apart from their hosts. In contrast, plant-associated microbes are frequently found in soils, where they can alter environmental microbial communities and generate feedbacks that influence plant community dynamics (i.e. Janzen-Connell effects; Van Der Heijden et al. 2008; Mangan et al. 2010). Overall, our results suggest that horizontal transmission links microbiomes among animal hosts but does not restructure environmental microbial communities. Therefore, host community dynamics may alter microbiomes in hosts while remaining decoupled from environmental microbial communities.

### 4.2 Effects of host identity, density, and diversity on microbiome diversity

Zooplankton Shannon diversity affected the alpha-diversity of host-associated microbiomes, and these effects differed among zooplankton taxa (Fig 3). We predicted that microbiome diversity would either increase through enhanced microbial transmission among heterospecifics or decline due to dilution of conspecific transmission (Fig 1B). We found asymmetric responses of microbiome diversity to host diversity among different zooplankton taxa (Fig 3). Microbiome alpha-diversity increased with zooplankton diversity in *Calanoida*, decreased in *Daphnia,* and was insensitive in *Cyclopoida*. These patterns suggest several potential mechanisms linking host and microbiome diversity.

First, the densities of different zooplankton hosts covaried with host diversity (Figs 2D – 2F). Rates of conspecific and heterospecific microbial transmission are likely density dependent. Unlike plant biodiversity experiments that maintain fixed richness treatments through weeding (Tilman & Downing 1994; Tilman et al. 1997; Hector et al. 1999), our diversity gradient emerged dynamically through temporal variation in the richness and relative abundance of hosts. Shifts in host abundance affect the transmission of microbes among host taxa and can influence whether host diversity amplifies or depresses microbiome diversity (Fig 1B). Prior work shows that increasing conspecific density can elevate microbiome richness (Burns et al. 2017) and that host population declines are associated with altered microbiome diversity (Pearman et al. 2024). Incompetent heterospecific hosts may act as “dead ends”, depressing local diversity when conspecifics are less abundant. The two most abundant host taxa, *Daphnia* and *Cyclopoida*, exhibited patterns consistent with dilution effects. *Daphnia* microbiome alpha diversity declined with increasing heterospecific host density (Fig 3C; Table 1). Because Daphnia density was negatively correlated with zooplankton Shannon diversity, more diverse host communities likely reduced opportunities for conspecific microbial exchange among *Daphnia*. Similarly, *Cyclopoida* microbiome richness declined as heterospecific density increased (Table 1; Table S8). In contrast, *Calanoida* microbiomes were insensitive to direct density effects (Table 1) and calanoids were comparatively rare within host communities (Fig 2C). Increasing host diversity may have amplified calanoid microbiome diversity through enhanced transmission among heterospecific hosts, consistent with our first hypothesis (Fig 1B). Overall, these results suggest that host diversity can both amplify and constrain microbiome diversity depending on the relative abundance and transmission dynamics of different host taxa and whether hosts are most abundant in high or low diversity communities.

### 4.3 Host densities regulate host-associated indicator abundance

Changes in host community structure may disproportionately affect microbes tightly connected to hosts. The relative abundance of *Calanoida* indicator microbes declined in calanoid copepods as the abundance of heterospecific hosts increased (Fig 4C), and *Daphnia* indicators increased in relative abundance in *Daphnia* as conspecifics became more abundant (Fig 4D). Despite representing few microbial families, these indicator taxa numerically dominated host microbiomes (Fig 4B), whereas most microbial diversity consisted of low-abundance zooplankton generalist and environmental-associated families (Fig 4A). Host filtering and/or selection of symbionts may explain this pattern, demonstrating that changes in the density of conspecific and heterospecific hosts alters the abundance of host-specialist microbes.

These results have implications for understanding the distribution of microbial symbionts that exhibit high host-affinity. Theory in disease ecology predicts that pathogen infection rates vary as a function of density-dependent transmission among hosts (Anderson & Mary 1991; Hochachka & Dhondt 2000). Our results suggest a similar dynamic in host-associated microbiomes. Microbes specialized to specific hosts increased in abundance with higher conspecific density or declined as heterospecific hosts became more abundant (Figs 4C & 4D). These patterns suggest that heterospecific hosts may act as “dead ends” for indicator taxa and potentially dilute their transmission among conspecifics. Importantly, shifts in the abundance of host-associated indicator taxa may have consequences for hosts. Previous work demonstrates that host-specific taxa can affect fitness and influence responses to environmental change (Jackrel et al. 2021; Fontaine et al. 2022). For example, *Flavobacteriaceae*, which were indicators of *Daphnia* in our study, have been implicated in local adaptation to environmental disturbance (Houwenhuyse et al. 2021). Thus, density-dependent changes in host communities not only restructure microbiomes but may alter host performance through changes in the distribution of key microbial taxa.

### 4.4 Host turnover drives turnover in host-associated microbiomes

Turnover in zooplankton community structure drove turnover in calanoid copepod and *Daphnia* microbiomes but not in *Cyclopoida* or tank water microbial communities (Fig 5A). Microbial communities in all sample types also exhibited temporal turnover, with the strongest effects observed in the environment (Fig 5B). This result suggests that free-living microbes are more temporally dynamic while hosts maintain a relatively consistent microbiome. While turnover in microbial communities is commonly attributed to environmental gradients (Bryant et al. 2008; Nemergut et al. 2013; Wall & Perreault et al. 2025), temporal dynamics (Lin et al. 2012; Martinović et al. 2021), and spatial scale (Martiny et al. 2011; Clark et al. 2021; Härer et al. 2025), our results show that changes in host communities themselves can shape microbial community structure. This suggests that shifts in macroscopic communities may alter patterns of horizontal transmission of microbes among hosts, linking host turnover to microbial community assembly. Importantly, these effects were not apparent in the environmental community, demonstrating that feedbacks between host and environmental microbiomes may be limited (Härer & Rennison 2023). These results suggest host community turnover may be a key, yet understudied, driver of beta diversity in host-associated microbiomes.

### 4.5 Conclusion

This study provides evidence that the structure of host communities can reshape microbiomes. We show that host diversity alters microbiome diversity, turnover, and the abundance of host-associated indicator microbes. Zooplankton diversity influenced microbiomes through shifts in the densities of individual host taxa that altered opportunities for microbial transmission among conspecific hosts. However, these effects did not extend to bacterioplankton, suggesting that feedbacks between host-associated and environmental microbial pools may be limited. Our results have implications for hosts because microbiome diversity can directly influence host health (Dickey et al. 2025; Corral López et al. 2026) and horizontally transmitted microbes can shape immunity (Koch & Schmid-Hempel 2011). Consequently, determining whether these microbiome shifts alter host fitness and scale up to influence population dynamics, coexistence, and ecosystem processes remains a critical next step.

