## Supporting Information for "Does host diversity beget microbiome diversity? Effects of zooplankton diversity and composition on environmental and host-associated microbes"

**This PDF file includes:**

Supplemental Methods

Supplemental Figures S1 – S3

Supplemental Tables S1 – S11

Supplemental References

### SUPPLEMENTAL METHODS

#### *Mesocosm Experimental Set-up*

During the summer season of 2022 (July – August), we assembled a field experiment to test the effects of host community diversity and density on host (zooplankton) and environmental (pond bacterioplankton) microbial communities. The experiment was performed outdoors in aquatic mesocosm tanks housed at the Sierra Nevada Aquatic Research Laboratory (SNARL). Prior to the start of the experiment, we established functioning aquatic ecosystems by filling twenty-four 1000 L mesocosm tanks with stream water. When filling tanks, we filtered water through a 63-$\mu\text{m}$ zooplankton net to prevent the establishment of any resident zooplankton from the stream. Next, tanks were fertilized with 264 $\mu\text{g/L}$ of nitrogen ($\text{NaNO}_3$) and 27 $\mu\text{g/L}$ of phosphorous ($\text{KH}_2\text{PO}_4$) (Kratina et al. 2012) and covered with shade cloth.

To establish host community treatments, we repeatedly collected zooplankton from Eastern Brook Lake (37.43153°N, 118.74262°W) using a 64-$\mu\text{m}$ mesh zooplankton net daily for two weeks. After collection each morning, zooplankton were transferred back to SNARL in cooled 2 L thermoses and living focal zooplankton taxa (calanoid copepods, cyclopoid copepods, and *Daphnia*) were separated into 1 L thermoses using a dissecting microscope. Next, we introduced zooplankton into mesocosms in 8 treatments replicated 3-times (no zooplankton, 3 single focal taxa treatments, 3 pairwise-focal taxa treatments, and a treatment with all three focal taxa). We repeatedly introduced zooplankton into treatments to ensure sufficient densities. After two weeks of stocking, we ceased community manipulations.

#### *DNA Library Preparation and Sequencing*

Zooplankton DNA was extracted with the MagMax Microbiome Ultra Nucleic Acid Extraction kit (ThermoFisher Scientific, USA) using the Kingfisher Apex System (ThermoFisher Scientific, USA). For bacterioplankton, we extracted DNA from 72 water filters (0.2-μm) using the Qiagen DNeasy PowerWater Kit (Qiagen, cat no. 14900-100-NF) according to manufacturer protocols. Blanks were used during extractions to identify potential contaminants. We quantified extracted DNA using the Qubit High Sensitivity DNA quantification kit (ThermoFisher Scientific, USA).

Following DNA extraction and quantification in house, 30 μL aliquots of samples were sent to the Argonne National Laboratory (ANL) for library preparation and 250 × 250 bp paired-end Illumina MiSeq sequencing (Caporaso et al. 2012). The V4 region of the 16S rRNA gene was amplified using modified primers that increase detection of aquatic microbes: 515F (5'-GTGYCAGCMGCCGCGGTAA-3') and 806R (5'-GGACTACNVGGGTWTCTAAT-3'). Illumina flowcell adaptor sequences were included on primers, and a 12 bp barcode sequence was attached to the forward primer to allow for pooling. PCR reactions (25 μL) contained 9.5 μL of MO BIO PCR Water (Certified DNA-Free), 12.5 μL of QuantaBio's AccuStart II PCR ToughMix (2x concentration), 1 μL Golay barcode tagged Forward Primer (5 μM concentration), 1 μL Reverse Primer (5 μM concentration), and 1 μL of template DNA. Thermocycler conditions consisted of 94°C for 3 minutes; 35 cycles of 94°C for 45 s, 50°C for 60 s, and 72°C for 90 s; and 72°C for 10 minutes. Post-PCR DNA was quantified using PicoGreen (Invitrogen) and pooled in equimolar volumes. Pool amplicons were bead cleaned (AMPure XP Beads; Beckman Coulter), quantified with Qubit (Invitrogen), and diluted before sequencing.

#### *Bioinformatic processing*

We ran demultiplexed fastq files through the DADA2 pipeline to identify ASVs (Callahan et al. 2016). After quality inspection and truncation, forward and reverse reads were merged and resolved to amplicon sequence variants (ASVs). Chimeric sequences were removed and taxonomy was assigned to ASVs using the 2021 SILVA 138.1 16S rRNA sequence database (Quast et al. 2013). Extraction controls were used to remove potential contaminants with the R package ‘decontam’ (Davis et al. 2018). Samples with fewer than 100 reads were excluded, and ASVs occurring in less than two samples or classified as chloroplasts or mitochondria were removed using the *phyloseq* package (McMurdie & Holmes 2013). A phylogenetic tree of ASVs was calculated using the *nj()* from the ‘ape’ R package (Paradis & Schliep 2019). To account for unequal sequencing depth, ASV tables were normalized to 7500 reads using scaling with ranked subsampling in the ‘SRS’ R package (Beule & Karlovsky 2020; Heidrich et al. 2021).

### SUPPLEMENTAL FIGURES

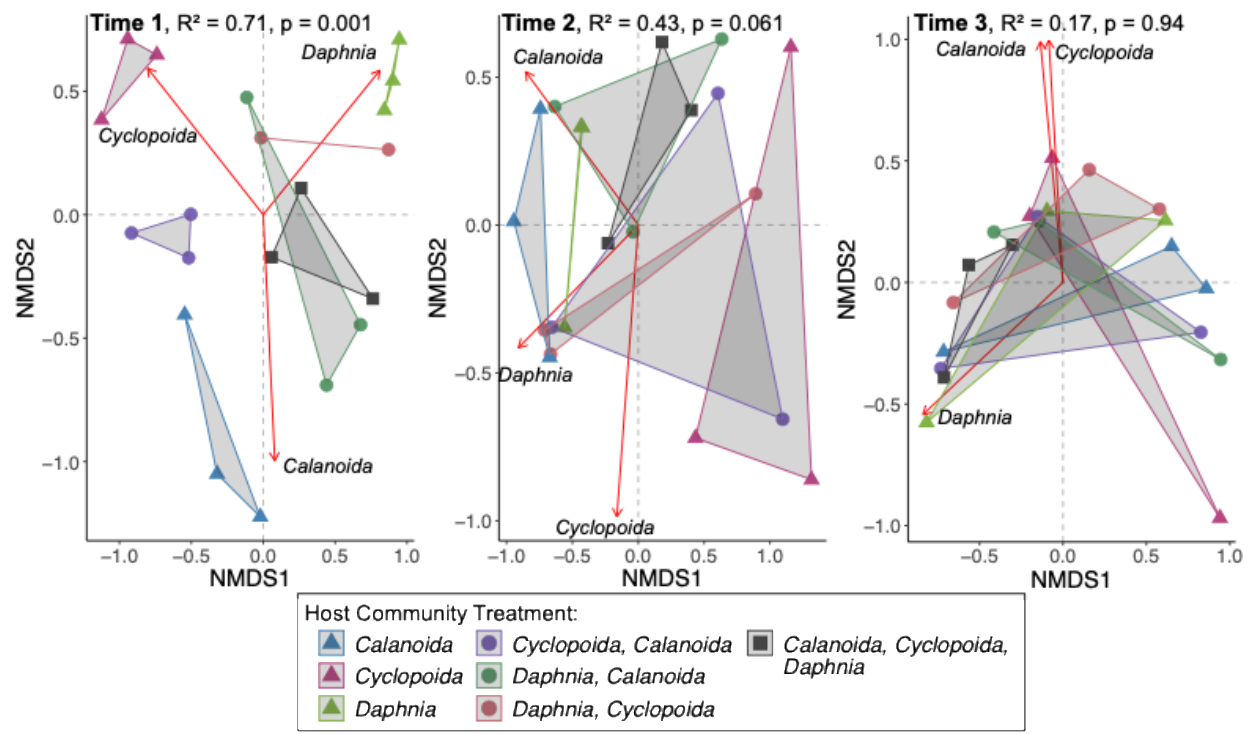

**Figure S1.** Non-metric multidimensional scaling (NMDS) depicting zooplankton community structure at each time point. Host community treatments are represented by color and richness treatment (1, 2, or 3 zooplankton taxa) are represented by shapes. $R^2$ and P-values represent the amount of variation explained and the significance of community treatment at each time point. Vectors correspond to zooplankton taxa whose abundances are correlated with NMDS axes.

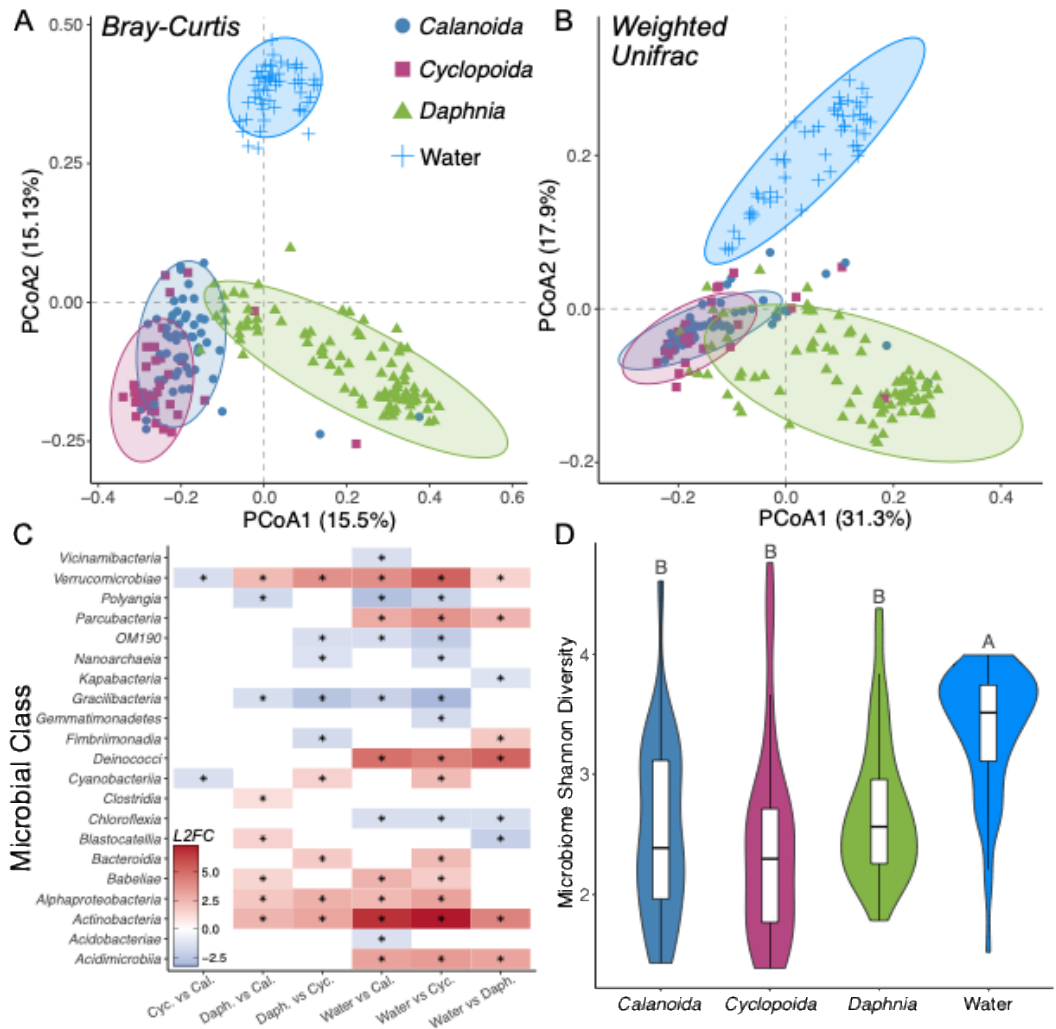

**Figure S2. (A, B)** Principle Coordinate Analysis (PCoA) plots using Bray-Curtis and Weighted-Unifrac measures of community dissimilarity across hosts (*Calanoida*, *Cyclopoida*, *Daphnia*) and the surrounding water column. Color and shape denote sample type. Ellipses represent the 95% confidence interval around the centroid of a given sample type. **(C)** Heatmap depicting the differential abundance (log2 fold differences in abundance) of microbial classes between sample types. Red plots indicate taxa enriched in the first term of a given pairwise comparison. **(D)** Violin plots of microbiome Shannon diversity across sample types. Letters indicate whether pairwise differences between groups were significantly different based on post-hoc tests ($p < 0.05$).

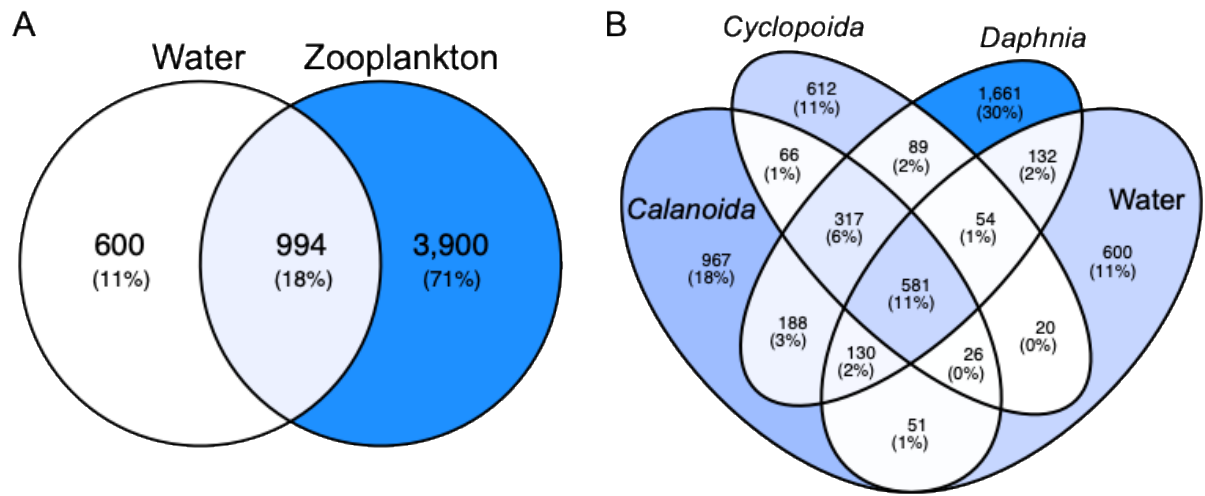

**Figure S3.** Venn diagram of the number and percentages of shared ASVs between **(A)** zooplankton hosts and water and **(B)** among all sample types.

### SUPPLEMENTAL TABLES:

**Table S1.** Results of linear mixed effects model evaluating the effects of time and richness treatment on zooplankton Shannon diversity.

| *Term* | *Estimate* | *SE* | *z* | *P* |
| --- | --- | --- | --- | --- |
| **Intercept (Time 1, Richness 1)** | 0.393 | 0.073 | 5.42 | **<0.001** |
| **Time Point** |  |  |  |  |
| Time 2 | 0.013 | 0.086 | 0.15 | 0.881 |
| Time 3 | 0.126 | 0.086 | 1.46 | 0.144 |
| **Richness Treatment** |  |  |  |  |
| Richness 2 | 0.282 | 0.105 | 2.67 | **0.008** |
| Richness 3 | 0.353 | 0.145 | 2.43 | **0.015** |
| **Time x Richness** |  |  |  |  |
| Time 2 × Richness 2 | -0.088 | 0.124 | -0.71 | 0.480 |
| Time 3 × Richness 2 | -0.262 | 0.124 | -2.11 | **0.035** |
| Time 2 × Richness 3 | -0.393 | 0.172 | -2.28 | **0.023** |
| Time 3 × Richness 3 | -0.722 | 0.172 | -4.19 | **<0.001** |

Estimates generated from linear mixed effects model with a Gaussian distribution. *Estimate* = coefficient estimate; *SE* = standard error of estimate; *z* = test statistic. Significant effects ($P < 0.05$) are in bold.

**Table S2.** Pairwise comparisons of zooplankton Shannon diversity in each richness treatment within each time point (Tukey-adjusted).

| *Time Point* | *Contrast* | *Estimate* | *SE* | *t.ratio* | *P* |
| --- | --- | --- | --- | --- | --- |
| Time Point 1 | Rich Treat 1 - Rich Treat 2 | -0.281 | 0.105 | -2.67 | **0.027** |
|  | Rich Treat 1 - Rich Treat 3 | -0.353 | 0.145 | -2.43 | **0.047** |
|  | Rich Treat 2 - Rich Treat 3 | -0.072 | 0.147 | -0.49 | 0.878 |
| Time Point 2 | Rich Treat 1 - Rich Treat 2 | -0.194 | 0.103 | -1.89 | 0.15 |
|  | Rich Treat 1 - Rich Treat 3 | 0.039 | 0.145 | 0.27 | 0.96 |
|  | Rich Treat 2 - Rich Treat 3 | 0.233 | 0.145 | 1.61 | 0.25 |
| Time Point 3 | Rich Treat 1 - Rich Treat 2 | -0.020 | 0.103 | -0.19 | 0.98 |
|  | Rich Treat 1 - Rich Treat 3 | 0.369 | 0.145 | 2.54 | **0.037** |
|  | Rich Treat 2 - Rich Treat 3 | 0.389 | 0.145 | 2.68 | **0.027** |

Pairwise difference estimates generated from post-hoc tukey tests. *Estimate* = coefficient estimate; *SE* = standard error of estimate; *t.ratio* = test statistic. Significant effects ($P < 0.05$) are in bold.

**Table S3.** Results of GLMM evaluating the effects of host identity and time on zooplankton densities.

| *Term* | *Estimate* | *SE* | *z* | *P* |
| --- | --- | --- | --- | --- |
| **Intercept (*Calanoida*, Time 1)** | 1.579 | 0.324 | 4.88 | **<0.001** |
| **Host (Time 1)** |  |  |  |  |
| *Cyclopoida* | 0.739 | 0.452 | 1.63 | 0.102 |
| *Daphnia* | 0.963 | 0.451 | 2.14 | **0.033** |
| **Time Point** |  |  |  |  |
| Time Point 2 | 0.478 | 0.443 | 1.08 | 0.282 |
| Time Point 3 | 0.156 | 0.437 | 0.36 | 0.722 |
| **Host × Time Point** |  |  |  |  |
| *Cyclopoida* × Time Point 2 | -0.633 | 0.622 | -1.02 | 0.309 |
| *Daphnia* × Time Point 2 | 0.948 | 0.618 | 1.53 | 0.125 |
| *Cyclopoida* × Time Point 3 | 0.52 | 0.609 | 0.85 | 0.400 |
| *Daphnia* × Time Point 3 | 2.40 | 0.607 | 3.95 | **<0.001** |

Estimates generated from generalized linear mixed effects model with a negative binomial distribution. *Estimate* = coefficient estimate; *SE* = standard error of estimate; *z* = test statistic. Significant effects ($P < 0.05$) are in bold.

**Table S4.** Pairwise comparisons of host densities within each time point (Tukey-adjusted).

| *Time Point* | *Contrast* | *Estimate* | *SE* | *z.ratio* | *P* |
| --- | --- | --- | --- | --- | --- |
| Time Point 1 | *Calanoida - Cyclopoida* | -0.738 | 0.452 | -1.63 | 0.231 |
|  | *Calanoida - Daphnia* | -0.963 | 0.453 | -2.14 | 0.083‡ |
|  | *Cyclopoida - Daphnia* | -0.224 | 0.445 | -0.50 | 0.870 |
| Time Point 2 | *Calanoida - Cyclopoida* | -0.105 | 0.428 | -0.25 | 0.968 |
|  | *Calanoida - Daphnia* | -1.91 | 0.423 | -4.52 | **<0.001** |
|  | *Cyclopoida - Daphnia* | -1.81 | 0.422 | -4.28 | **0.001** |
| Time Point 3 | *Calanoida - Cyclopoida* | -1.23 | 0.409 | -3.07 | **0.006** |
|  | *Calanoida - Daphnia* | -3.36 | 0.406 | -8.27 | **<0.001** |
|  | *Cyclopoida - Daphnia* | -2.11 | 0.400 | -5.27 | **<0.001** |

Pairwise difference estimates generated from post-hoc tukey tests. *Estimate* = coefficient estimate; *SE* = standard error of estimate; *z.ratio* = test statistic. Significant effects ($P < 0.05$) are in bold and marginal effects ($P < 0.10$) are marked by ‡.

**Table S5.** PERMANOVA results testing the effects of community treatment, richness treatment, and time point on zooplankton community composition.

| ***Model 1*** | *df* | *SS* | *R<sup>2</sup>* | *F* | *Pr(>F)* |
| --- | --- | --- | --- | --- | --- |
| Time Point | 2 | 2.61 | 0.16 | 8.94 | **0.001** |
| Community Treatment | 7 | 4.41 | 0.26 | 4.31 | **0.001** |
| Time Point × Community Treatment | 13 | 3.39 | 0.20 | 1.79 | **0.041** |
| Residual | 43 | 6.28 | 0.38 |  |  |
| Total | 65 | 16.7 | 1.00 |  |  |

| ***Model 2*** | *df* | *SS* | *R<sup>2</sup>* | *F* | *Pr(>F)* |
| --- | --- | --- | --- | --- | --- |
| Time Point | 2 | 2.61 | 0.16 | 6.41 | **0.001** |
| Richness Treatment | 1 | 0.73 | 0.04 | 3.58 | **0.001** |
| Time Point × Richness Treatment | 2 | 1.14 | 0.07 | 2.80 | **0.039** |
| Residual | 60 |  | 0.73 |  |  |
| Total | 239 |  | 1.00 |  |  |

PERMANOVA tables generated from 999 permutations; *df* = degrees of freedom; *SS* = sum of squares. Significant effects ($P < 0.05$) are in bold.

**Table S6.** PERMANOVA output table evaluating the effects of sample type and time on microbiome community composition

| ***Bray-Curtis*** | *df* | *SS* | *R<sup>2</sup>* | *F* | *Pr(>F)* |
| --- | --- | --- | --- | --- | --- |
| Sample Type | 3 | 21.9 | 0.30 | 37.7 | **0.001** |
| Time Point | 2 | 5.9 | 0.08 | 15.3 | **0.001** |
| Residual | 234 | 45.3 | 0.61 |  |  |
| Total | 239 | 73.9 | 1.00 |  |  |
| ***Weighted-Unifrac*** |  |  |  |  |  |
| Sample Type | 3 | 7.23 | 0.37 | 52.3 | **0.001** |
| Time Point | 2 | 1.38 | 0.07 | 15.0 | **0.001** |
| Residual | 234 | 10.8 | 0.55 |  |  |
| Total | 239 | 19.6 | 1.00 |  |  |

PERMANOVA tables generated from 999 permutations; *df* = degrees of freedom; *SS* = sum of squares. Significant effects ($P < 0.05$) are in bold.

**Table S7.** PERMANOVA output table for the effects of host Shannon diversity, density, and time on microbiome composition. The full model contained sample type

| ***Bray-Curtis*** | *df* | *SS* | *R<sup>2</sup>* | *F* | *Pr(>F)* |
| --- | --- | --- | --- | --- | --- |
| Time Point | 2 | 4.36 | 0.06 | 11.7 | **0.001** |
| Sample Type × Host Shannon | 3 | 0.97 | 0.01 | 1.75 | **0.044** |
| Sample Type × Total Host Density | 3 | 1.44 | 0.02 | 2.59 | **0.001** |
| Residual | 226 | 42.1 | 0.56 |  |  |
| Total | 239 | 73.9 | 1.00 |  |  |
| ***Weighted-Unifrac*** |  |  |  |  |  |
| Time Point | 2 | 0.81 | 0.04 | 9.50 | **0.001** |
| Sample Type × Host Shannon | 3 | 0.22 | 0.01 | 1.73 | 0.108 |
| Sample Type × Total Host Density | 3 | 0.56 | 0.03 | 4.34 | **0.001** |
| Residual | 226 | 9.69 | 0.49 |  |  |
| Total | 239 | 19.6 | 1.00 |  |  |

PERMANOVA tables generated from 999 permutations; *df* = *degrees of freedom*; *SS* = *sum of squares*. Significant effects ($P < 0.05$) are in bold.

**Table S8.** Output tables from generalized linear mixed effects models predicting microbial alpha-diversity as a function of zooplankton diversity or zooplankton density.

|  | *Alpha Metric* | *Model* | *Term* | *Estimate* | *SE* | *P* | *AICc* | *ΔAICc* |
| --- | --- | --- | --- | --- | --- | --- | --- | --- |
| ***Calanoida*** | Shannon | Diversity | Host Shannon | 0.131 | 0.038 | **<0.001** | 119 | 0 |
|  | Shannon | Density | Con. Density | 0.018 | 0.039 | 0.650 | 128.9 | 9.9 |
|  | Shannon |  | Het. Density | -0.013 | 0.069 | 0.850 | 128.9 |  |
|  | Richness | Diversity | Host Shannon | 0.144 | 0.08 | 0.07‡ | 656.1 | 0 |
|  | Richness | Density | Con. Density | 0.056 | 0.069 | 0.410 | 660.9 | 4.8 |
|  | Richness |  | Het. Density | -0.009 | 0.068 | 0.900 | 660.9 |  |
|  | Pielou | Diversity | Host Shannon | 0.193 | 0.064 | **0.002** | -97.2 | 0 |
|  | Pielou | Density | Con. Density | 0.019 | 0.055 | 0.730 | -86 | 11.2 |
|  | Pielou |  | Het. Density | -0.006 | 0.096 | 0.950 | -86 |  |
| ***Cyclopoida*** | Shannon | Diversity | Host Shannon | -0.012 | 0.043 | 0.780 | 90.4 | 0.1 |
|  | Shannon | Density | Con. Density | 0.017 | 0.044 | 0.690 | 90.3 | 0 |
|  | Shannon |  | Het. Density | -0.089 | 0.052 | 0.09‡ | 90.3 |  |
|  | Richness | Diversity | Host Shannon | -0.089 | 0.094 | 0.340 | 473.9 | 3.9 |
|  | Richness | Density | Con. Density | 0.133 | 0.091 | 0.140 | 470 | 0 |
|  | Richness |  | Het. Density | -0.217 | 0.068 | **0.001** | 470 |  |
|  | Pielou | Diversity | Host Shannon | -0.017 | 0.069 | 0.810 | -49.5 | 0 |
|  | Pielou | Density | Con. Density | -0.009 | 0.077 | 0.900 | -47.5 | 2 |
|  | Pielou |  | Het. Density | -0.083 | 0.091 | 0.360 | -47.5 |  |
| ***Daphnia*** | Shannon | Diversity | Host Shannon | -0.066 | 0.022 | **0.003** | 121.9 | 0 |
|  | Shannon | Density | Con. Density | 0.016 | 0.047 | 0.730 | 123.5 | 1.6 |
|  | Shannon |  | Het. Density | -0.069 | 0.022 | **0.002** | 123.5 |  |
|  | Richness | Diversity | Host Shannon | -0.126 | 0.044 | **0.004** | 1014.3 | 0 |
|  | Richness | Density | Con. Density | 0.182 | 0.101 | 0.07‡ | 1017.7 | 0.4 |
|  | Richness |  | Het. Density | -0.094 | 0.045 | **0.037** | 1017.7 |  |
|  | Pielou | Diversity | Host Shannon | -0.08 | 0.033 | **0.015** | -219.2 | 0.7 |
|  | Pielou | Density | Con. Density | -0.032 | 0.063 | 0.610 | -220.9 | 0 |
|  | Pielou |  | Het. Density | -0.103 | 0.032 | **0.001** | -220.9 |  |
| **Water** | Shannon | Diversity | Host Shannon | 0.027 | 0.018 | 0.130 | 81.1 | 0 |
|  | Shannon | Density | Tot. Host Density | -0.007 | 0.021 | 0.760 | 83.3 | 2.2 |
|  | Richness | Diversity | Host Shannon | 0.038 | 0.028 | 0.180 | 565.8 | 0 |
|  | Richness | Density | Tot. Host Density | 0.003 | 0.029 | 0.930 | 567.5 | 1.7 |
|  | Pielou | Diversity | Host Shannon | 0.046 | 0.031 | 0.140 | -139.3 | 0 |
|  | Pielou | Density | Tot. Host Density | -0.029 | 0.039 | 0.460 | -137.7 | 1.6 |

Estimates generated from generalized linear mixed effects models. *Estimate* = coefficient estimate; *SE* = standard error of estimate. Significant predictors ($P < 0.05$) are in bold and marginal predictors ($P < 0.10$) are marked by ‡.

**Table S9.** Results of Indicator Species Analysis.

|  | *Mean Relative Abundance* | *Indicator Value* | *P* |
| --- | --- | --- | --- |
| ***Calanoida*** |  |  |  |
| *Comamonadaceae* | 0.57368333 | 0.63465604 | **0.001** |
| *Polyangiaceae* | 0.00117143 | 0.2999839 | **0.001** |
| *Myxococcaceae* | 0.00019762 | 0.18651444 | **0.027** |
| *Sandaracinaceae* | 0.00012143 | 0.17808009 | **0.036** |
| ***Cyclopoida*** |  |  |  |
| *Aeromonadaceae* | 0.41077398 | 0.73566366 | **0.001** |
| *Corynebacteriaceae* | 0.00481626 | 0.24827135 | **0.001** |
| *Rickettsiaceae* | 0.00333984 | 0.25992982 | **0.001** |
| *Staphylococcaceae* | 0.00323902 | 0.21246269 | **0.005** |
| *Streptococcaceae* | 0.00040976 | 0.15002716 | **0.047** |
| *Vibrionaceae* | 0.00037724 | 0.16548494 | **0.032** |
| *Halomonadaceae* | 0.00025366 | 0.18931869 | **0.003** |
| *Rhodocyclaceae* | 0.00021463 | 0.24341701 | **0.001** |
| *Candidatus Jidaibacter* | 0.00020813 | 0.15293103 | **0.004** |
| *Hydrogenophilaceae* | 0.00012358 | 0.13822809 | **0.044** |
| *Micavibrionaceae* | 8.13E-05 | 0.16897319 | **0.046** |
| *Anaerovoracaceae* | 5.85E-05 | 0.17850847 | **0.014** |
| *Rhodothermaceae* | 4.23E-05 | 0.14467467 | **0.026** |
| ***Daphnia*** |  |  |  |
| *Candidatus Hepatincola* | 0.24776522 | 0.73628019 | **0.001** |
| *T34* | 0.15674783 | 0.71245183 | **0.001** |
| *Flavobacteriaceae* | 0.13609565 | 0.2779087 | **0.001** |
| *Rhodobacteraceae* | 0.0696 | 0.2329376 | **0.002** |
| *Amoebophilaceae* | 0.00141159 | 0.27155246 | **0.001** |
| *Reyranellaceae* | 0.00038551 | 0.19793487 | **0.017** |
| ***Water*** |  |  |  |
| *Sporichthyaceae* | 0.11241975 | 0.69586572 | **0.001** |
| *Burkholderiaceae* | 0.02860741 | 0.6552811 | **0.001** |
| *Microbacteriaceae* | 0.01061728 | 0.57665635 | **0.001** |
| *Chitinophagaceae* | 0.03462222 | 0.55061924 | **0.001** |
| *Alcaligenaceae* | 0.01483457 | 0.55037146 | **0.001** |
| *Sphingomonadaceae* | 0.04757037 | 0.54572461 | **0.001** |
| *Clade III* | 0.03981975 | 0.46025975 | **0.001** |
| *Mycobacteriaceae* | 0.01350617 | 0.43562092 | **0.001** |
| *Pseudomonadaceae* | 0.03731605 | 0.431818 | **0.001** |
| *Moraxellaceae* | 0.06389877 | 0.43134475 | **0.001** |
| *Cyclobacteriaceae* | 0.01087407 | 0.42067289 | **0.001** |
| *Methylophilaceae* | 0.00276543 | 0.41855085 | **0.001** |
| *Deinococcaceae* | 0.08820741 | 0.40936431 | **0.001** |
| *Rhizobiaceae* | 0.01824938 | 0.39683911 | **0.001** |
| *NS11-12 marine group* | 0.01473827 | 0.39501625 | **0.001** |

Mean relative abundances were calculated within each sample type. The top fifteen water-associated indicator families are shown. Indicator and P-values were derived from the indicator species analysis.

**Table S10.** Output table from generalized linear mixed effects models predicting the summed relative abundance of microbial indicators in focal hosts as a function of conspecific and heterospecific density.

| *Calanoida* Indicator Rel. Abundance | *Estimate* | *SE* | *z-value* | *P* | *R*<sup>2</sup><sub>*m*</sub> |
| --- | --- | --- | --- | --- | --- |
| Intercept | 0.14 | 0.20 | 0.68 | 0.50 | 0.35 |
| *log* Conspecific Density | 0.16 | 0.09 | 1.76 | 0.08‡ |  |
| *log* Heterospecific (*Daphnia* + *Cylopoida*) Density | -0.47 | 0.11 | -4.44 | **<0.001** |  |
| *Cyclopoida* Indicator Rel. Abundance |  |  |  |  |  |
| Intercept | -0.36 | 0.18 | -2.01 | 0.04 | 0.10 |
| *log* Conspecific Density | 0.16 | 0.34 | 0.45 | 0.65 |  |
| *log* Heterospecific (*Daphnia* + *Calanoida*) Density | -0.31 | 0.32 | -0.97 | 0.33 |  |
| *Daphnia* Indicator Rel. Abundance |  |  |  |  |  |
| Intercept | 0.13 | 0.48 | 0.28 | 0.78 | 0.08 |
| *log Daphnia* Density | 0.29 | 0.16 | 1.8 | 0.06‡ |  |
| *log* Heterospecific (*Calanoida* + *Cylopoida*) Density | 0.09 | 0.08 | 1.1 | 0.26 |  |

Estimates generated from generalized linear mixed effects models fit with a beta distribution. *Estimate* = coefficient estimate; *SE* = standard error of estimate, *R*<sup>2</sup><sub>*m*</sub> = variance explained by model fixed effects. Significant predictors (*P* < 0.05) are in bold and marginal predictors are marked by ‡ (*P* < 0.10).

**Table S11.** Parameter estimates from Bayesian generalized linear mixed effects model predicting microbial beta-diversity as a function of time (difference in days) and host beta-diversity.

| ***Microbial Turnover (Bray-Curtis)*** | $\beta$ | *Est.Err* | *95% CrI* | $R^2$ |
| --- | --- | --- | --- | --- |
| Intercept (*Calanoida*) | 0.71 | 0.08 | [0.56, 0.86] | 1.00 |
| *Cyclopoida* (vs *Calanoida*) | -0.18 | 0.12 | [-0.42, 0.06] | 1.00 |
| *Daphnia* (vs *Calanoida*) | -0.20 | 0.10 | [-0.39, 0.00] | 1.00 |
| Water (vs *Calanoida*) | 0.11 | 0.11 | [-0.11, 0.32] | 1.00 |
| Days (*Calanoida* Slope) | **0.13** | **0.01** | **[0.11, 0.15]** | 1.00 |
| Host Beta (*Calanoida* Slope) | **0.09** | **0.01** | **[0.07, 0.12]** | 1.00 |
| *Cyclopoida* × Days | **0.01** | **0.02** | **[-0.03, 0.05]** | 1.00 |
| *Daphnia* × Days | **0.11** | **0.01** | **[0.08, 0.14]** | 1.00 |
| Water × Days | **0.17** | **0.02** | **[0.14, 0.20]** | 1.00 |
| *Cyclopoida* × Host Beta | -0.07 | 0.02 | [-0.11, -0.03] | 1.00 |
| *Daphnia* × Host Beta | **-0.01** | **0.01** | **[-0.04, 0.01]** | 1.00 |
| Water × Host Beta | -0.10 | 0.02 | [-0.14, -0.07] | 1.00 |

Posterior estimates ($\beta$) of fixed effects on the logit scale from the Bayesian GLMM modeling microbial turnover (Bray-Curtis distance) as a function of host turnover (Bray-Curtis) and time (difference in days). The model was fit using a beta distribution with a logit link and interaction terms represent differences in slopes relative to *Calanoida*. Continuous predictors were z-standardized (mean = 0, SD = 1). Coefficients with credible effects (95% credible interval excludes zero) are in bold.

Improved bacterial 16S rRNA gene (V4 and V4-5) and fungal internal transcribed spacer marker gene primers for microbial community surveys. *mSystems*, 1(1).

https://doi.org/10.1128/mSystems.00009-15
